# Two light sensors decode moonlight versus sunlight to adjust a plastic circadian/circalunidian clock to moon phase

**DOI:** 10.1101/2021.04.16.440114

**Authors:** Martin Zurl, Birgit Poehn, Dirk Rieger, Shruthi Krishnan, Dunja Rokvic, Vinoth Babu Veedin Rajan, Elliot Gerrard, Matthias Schlichting, Lukas Orel, Robert J. Lucas, Eva Wolf, Charlotte Helfrich-Förster, Florian Raible, Kristin Tessmar-Raible

## Abstract

Many species synchronize their physiology and behavior to specific hours. It is commonly assumed that sunlight acts as the main entrainment signal for ~24h clocks. However, the moon provides similarly regular time information, and increasingly studies report correlations between diel behavior and lunidian cycles. Yet, mechanistic insight into the possible influences of the moon on ~24hr timers is scarce.

We studied *Platynereis dumerilii* and uncover that the moon, besides its role in monthly timing, also schedules the exact hour of nocturnal swarming onset to the nights’ darkest times. Moonlight adjusts a plastic clock, exhibiting <24h (moonlit) or >24h (no moon) periodicity. Abundance, light sensitivity, and genetic requirement indicate *Platynereis* r-Opsin1 as receptor to determine moonrise, while the cryptochrome L-Cry is required to discriminate between moon- and sunlight valence. Comparative experiments in *Drosophila* suggest that Cryptochrome’s requirement for light valence interpretation is conserved. Its exact biochemical properties differ, however, between species with dissimilar timing ecology.

Our work advances the molecular understanding of lunar impact on fundamental rhythmic processes, including those of marine mass spawners endangered by anthropogenic change.

## Main text

### A moonlight-sensitive clock times swarming behavior

*Platynereis dumerilii* reproduces by nocturnal mass spawning, with sexually mature males and females synchronously raising from seagrass to the water surface (Fig. 1A) during the night (*1*). While it is well established that this spawning is synchronized to specific nights of the month by a circalunar oscillator (refs. (*2*, *3*) and accompanying manuscript by Poehn, Krishnan et al), we reasoned that it should further increase reproductive success if worms synchronized the onset of swarming behavior also to specific hours during those nights. In fact, such an interconnection of different timing systems is well established for polychaete relatives like the palolo worms (*4*) and fireworms (*Odontosyllis*) (*5*).

**Figure 1.**
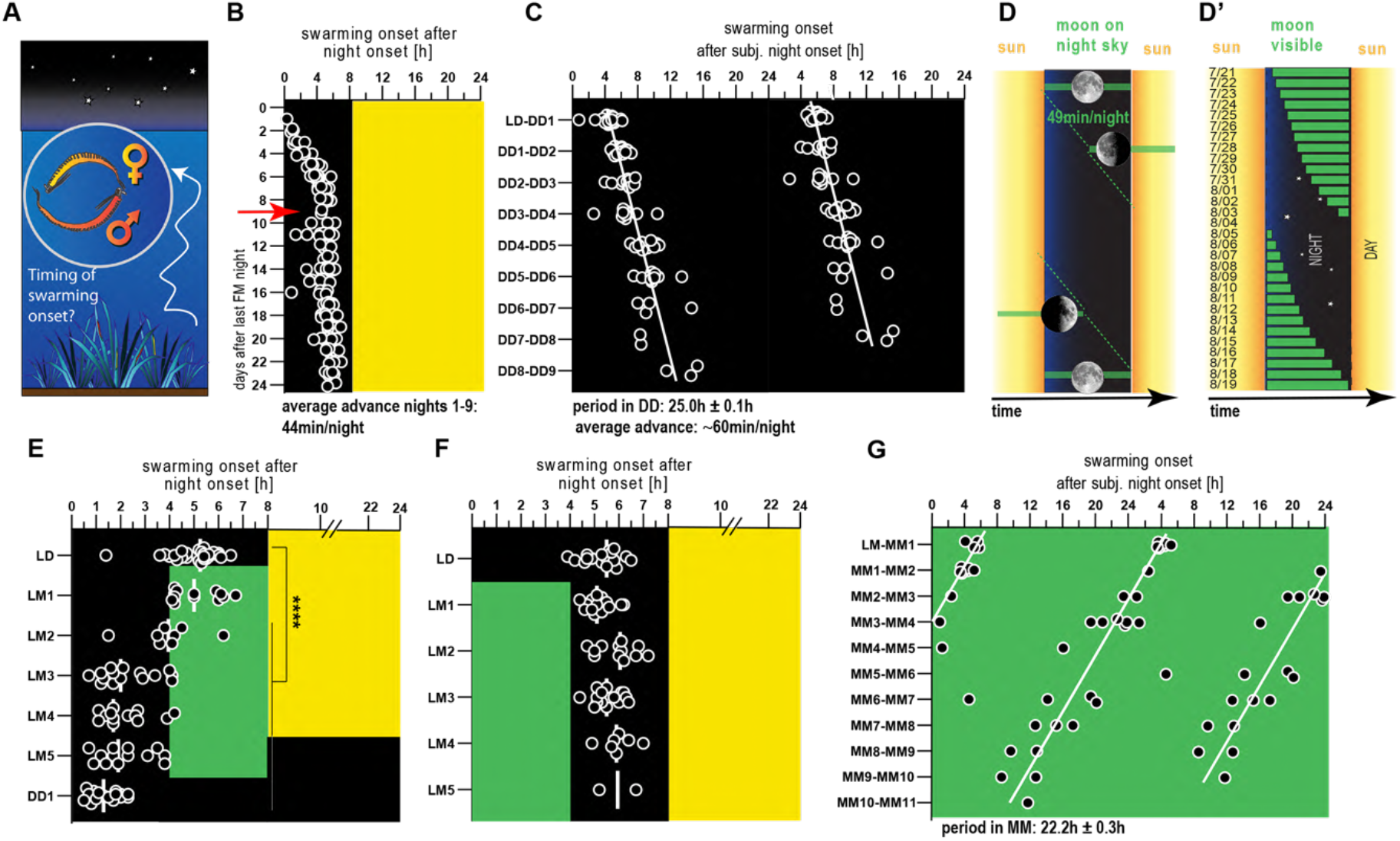
A moonlight sensitive plastic circadian/circalunidian clock (PCC) times swarming onset to darkness. **(A)** Schematized swarming behavior of *Platynereis dumerilii*. **(B)** Swarming onset of individual, separated worms across different days of an artificial lunar month, were worms receive 8 nights continuous nocturnal light (=full moon: FM) every month in addition to a 16h:8h LD cycle (for details see (*6*), accompanying manuscript Poehn, Krishnan et al). Red arrow in indicates days relative to the circalunar cycle from which onwards worms were used for all subsequent experiments (except Fig.2 F and G). **(C,G)** Swarming onset of worms released into constant darkness (DD; D) or constant moonlight (MM; G). Data are double-plotted for better visualization. White lines are linear regression lines. Period lengths were calculated based on the slope of the regression line ± the 95% CI of the slope. **(D,D’)** Schemes illustrating moon rise and set times in a simplified averaged model (D) and the natural situation (Bay of Naples, July/August 1929) (D’). See: https://www.timeanddate.com/moon/italy/naples **(E,F)** Swarming onset of worms subjected to naturalistic moonlight during the second (E) or first (F) half of the night. black: no light, yellow: naturalistic sunlight, green: naturalistic moonlight.

This prompted us to investigate if *Platynereis dumerilii* also exhibits preferred hours of spawning. We placed maturing, monthly (circalunar) entrained *Platynereis dumerilii* adults (*3*) in individual wells of our automated behavioral recording device (*7*). As swarming is accompanied by a burst of swimming activity (“nuptial dance”), analysis by automated video tracking allowed us to systematically deduce the time of swarming onset with respect to the daylight/darkness (LD:16:8h) cycle (fig. S1A,B Supplementary Video 1). Analyses of 139 individuals revealed that swarming onset across the culture was indeed synchronized to a ~1-2hr window during the night (Fig. 1B). (Note that we selected about equal numbers of spawning worms/night. Therefore, the monthly spawning synchronization is invisible.) The precise time point depended on the time since the last artificial “full moon” (FM) night (Fig. 1B), which is provided to entrain the worms’ monthly oscillator (*3*). In nights directly following the last “full moon” night, animals started the characteristic swarming behavior directly following night onset. This onset of swarming gradually shifted by app. 44min/night within the first 8 nights (Fig.1B: days preceding the red arrow). For the remaining lunar month, the time of swarming onset remained unaltered at ~5 h after night onset (Fig. 1B, fig. S1B). To assess whether this synchronization was driven by an endogenous oscillator, we next monitored swarming onset in worms that were kept in constant darkness for several days. Under these dark-dark (DD) conditions, swarming was still synchronously initiated, with an average delay of ~1h ± 0,1h per day (Fig 1C). This established that the specific hour of nocturnal swarming onset is controlled by an endogenous clock.

The time advance of about 44min within the first 8 nights after full moon is reminiscent of the average delay of the rise of the waning moon (~ 49min/night, Fig. 1D). This apparent delay of moon rise time relative to sunset is caused by the period difference of the daily solar cycle (24h) and the lunidian cycle (24.8h; the average timespan between two successive moon rises) (Fig. 1D). The latter matches the period length of the endogenous clock (~25h) controlling swarming onset under DD conditions (compare Fig. 1C,D). The combination of these facts let us speculate that this timing system could help to synchronize *Platynereis* swarming onset to the darkest hours of the night, but would require the moon for entrainment. Furthermore, the exact change of moon rise relative to sunset is not always exactly ~49min/night, but varies under natural conditions (Fig. 1D’), making an additional adjustment by moonlight likely advantageous. We thus next studied if the endogenous clock was sensitive to moonlight for its exact entrainment. To mimic moonlight and sunlight under laboratory conditions, we complemented available surface measurements (*8*) by analyzing systematic light measurements at a natural habitat of *Platynereis* (fig. S2A), which guided the design of “naturalistic sunlight” and “naturalistic moonlight” illumination devices (fig. S2B, see also accompanying manuscript Poehn, Krishnan et al, and ref. (*7*)).

We next exposed animals (>= 9 days after the end of the monthly nocturnal light stimulus, see red arrow Fig. 1B) to “naturalistic moonlight” (fig. S2B) provided during the second half of the night for 5 consecutive nights (Fig. 1E, LM1-5). In response to this light regime mimicking “waning moon”, worms shifted their swarming onset gradually into the dark portion of these “moonlit” nights (Fig. 1E). The advanced swarming onset caused by the “waning moonlight regime” persisted when worms were subsequently released into constant darkness (Fig. 1E: DD1), arguing that this shift was caused by an impact of moonlight on the endogenous clock, rather than being an acute masking effect (i.e. direct response to light). Consistent with timing the dark portion of the night, the same “naturalistic moonlight” provided during the first half of the night (mimicking times of waxing moon) did not impact on the worms’ hourly timing (Fig. 1F). Finally, under a constant “naturalistic moonlight” (MM) regime, spawning onset remained synchronized, but occurred with a markedly decreased period length of ~22.2h ± 0.4h, compared to DD conditions (Fig. 1C vs. G).

Taken together, these results suggest the existence of a plastic oscillator system that regulates nocturnal swarming onset, whose period is modulated by naturalistic moonlight. This results in a swarming preference during the dark portion of the night, consistent with natural observations. We refer to this clock as plastic circadian/circalunidian clock (PCC clock).

### L-Cry is required to correctly interpret sun– and moonlight

In order to understand how (naturalistic) sun– and moonlight are sensed and distinguished by this system, we next sought to identify photoreceptor(s) relevant for the light impact on the PCC clock. One candidate receptor of particular interest was *Platynereis* L-Cryptochrome (L-Cry), whose distant homolog Cry2 in the coral *Acropora* has been speculated to mediate moonlight sensation based on expression changes (*9*).

To assess if L-Cry is relevant for light input into the PCC clock, we analyzed a *Platynereis l-cry* loss-of-function strain (see accompanying manuscript Poehn, Krishnan et al). When exposed to constant darkness, *l-cry^−/−^* individuals still exhibited rhythmic initiation of swarming onset, with a period length (24.6h ± 0.3h) indistinguishable from wildtypes (Fig. 2A). This indicates that L-Cry is not required for the endogenous oscillation of the PCC clock.

**Figure 2.**
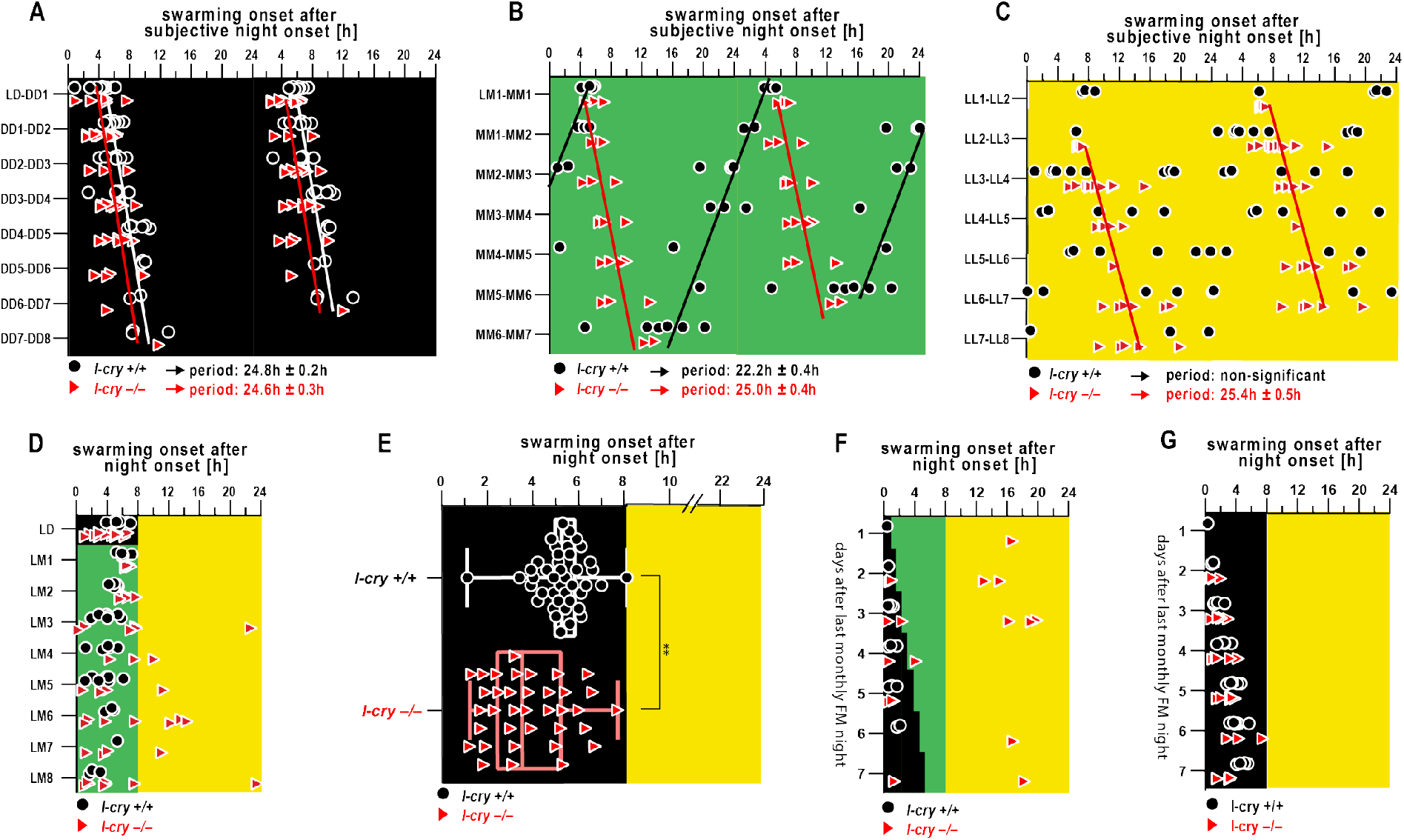
*Platynereis* L-Cry enables the PCC to distinguish sun- versus moonlight. **(A-E)** Swarming onset of *l-cry* mutants (red triangles) and wildtypes (black circles) entrained to 16:8h LD cycles subsequently released into **(A)** constant darkness (DD), **(B**) constant naturalistic moonlight (MM), or **(C)** constant naturalistic sunlight (LL), or **(D)** subjected to alternations of naturalistic sunlight during the day and moonlight during the night (LM) or **(E)** maintained under 16:8h LD cycles (**p=0.004, F-test to test if the variances in the two groups are significantly different). Data in A-C are double-plotted. Black and red lines indicate linear regression lines of wildtype and *l-cry^−/−^* mutants, respectively. The period length was calculated based on the slope of the regression line ± the 95% CI of the slope. **(F)** Swarming onset of *l-cry* mutants and wildtypes assessed directly after the monthly nocturnal full moon (FM) light stimulus and kept either under LD cycles (F) or with an additional waning moonlight regime (G).

To probe for roles of L-Cry in mediating light input into the PCC clock, we next investigated spawning rhythmicity in *l-cry* mutants exposed to constant “naturalistic moonlight” (MM) or “naturalistic sun light” (LL). Under both conditions, *l-cry* mutants exhibited a synchronized swarming onset, with period lengths (MM: 25h ± 0.4h; Fig. 2B; LL: 25.4h ± 0.5h. Fig. 2C) highly reminiscent of the period of wildtype in DD conditions (Fig. 2A). In contrast, wildtype siblings shortened their period (MM) or became arrhythmic (LL), respectively (Fig. 2B,C). These clear differences between wildtype and mutants let us conclude that L-Cry is relevant for the conveying naturalistic sun- and moonlight information to the PCC clock.

The absent adjustment of the PCC clock in *l-cry^−/−^* individuals to respond to light could be explained by a general reduction in light sensitivity. Alternatively, these findings are compatible with a role of L-Cry in distinguishing moon– and sunlight, as L-Cry enables the PCC clock to respond differently to the two light conditions. To discriminate between the two possibilities, we exposed *l-cry* mutants to a day/night regime of 16h:8h, where they were exposed to “naturalistic sunlight” during the day, and “naturalistic moonlight” during the night (LM) (Fig 2D). Unlike wildtype animals, that restricted swarming onset strictly to nocturnal hours, *l-cry* mutants exhibited aberrant swarming onset. Starting with 3 days of the LM regime, around a quarter of the recorded animals initiated swarming during the day, a phenomenon never observed for wildtype animals (Fig. 2D). In contrast, in LD conditions all *l-cry* mutants restricted swarming onset to the night, albeit less synchronized than wildtype, (Fig. 2D: LD,Fig.2E), indicating that the shifted timing into the day was caused by the naturalistic moonlight stimulus. The abnormal swarming onset of *l-cry^−/−^* animals was also observed in a light regime in which a staggered, artificial waning moonlight regime (fig. S2C) was provided directly after the end of the standard monthly culture FM stimulus, more closely mimicking the natural timing under which swarming is observed (Fig. 2F, compare Fig. 1D,D’) compared to the identical time and light regime lacking the waning moon stimulus (Fig.2G). Overall, this suggests that the *l-cry* mutation does not simply render worms less sensitive to moonlight, but that L-Cry is required to correctly interpret naturalistic moonlight versus sunlight stimuli.

### Subcellular localization and stability of L-Cry supports distinct signaling under moonlight and sunlight conditions

In the common view based on the work in *Drosophila melanogaster*, the fly homolog of L-Cry – dCry – undergoes light dependent binding to Timeless, which leads to the degradation of both Timeless and dCry, by this resetting the flies’ circadian clock upon light input (reviewed in ref. (*10*)). This binary signaling model is difficult to reconcile with our finding that *Platynereis* L-Cry is relevant for distinguishing between different light valences in the context of circadian/circalunidian timing.

We therefore tested if L-Cry protein in the worm exhibited any differences when animals were exposed to naturalistic sun–or moonlight under conditions relevant for the above behavioral paradigms. We made use of a *Pdu-*L-Cry-specific antibody (for antibody generation and validation see accompanying manuscript Poehn, Krishnan et al). We first assessed L-Cry abundance in head extracts of animals sampled at the midpoint of the subjective night (at new moon: NM), after 4h of darkness or exposure to either naturalistic sun- or moonlight (Fig. 3A, CT20, red arrows). As expected by the canonical *Drosophila* model and consistent with our previous analyses in S2 cells (*3*), naturalistic sunlight led to a significant reduction of L-Cry compared to heads sampled from animals maintained in darkness (Fig. 3B,C). In contrast, the levels of L-Cry protein in the heads of naturalistic moonlight-exposed animals was indistinguishable from dark levels (Fig. 3B,C).

**Figure 3.**
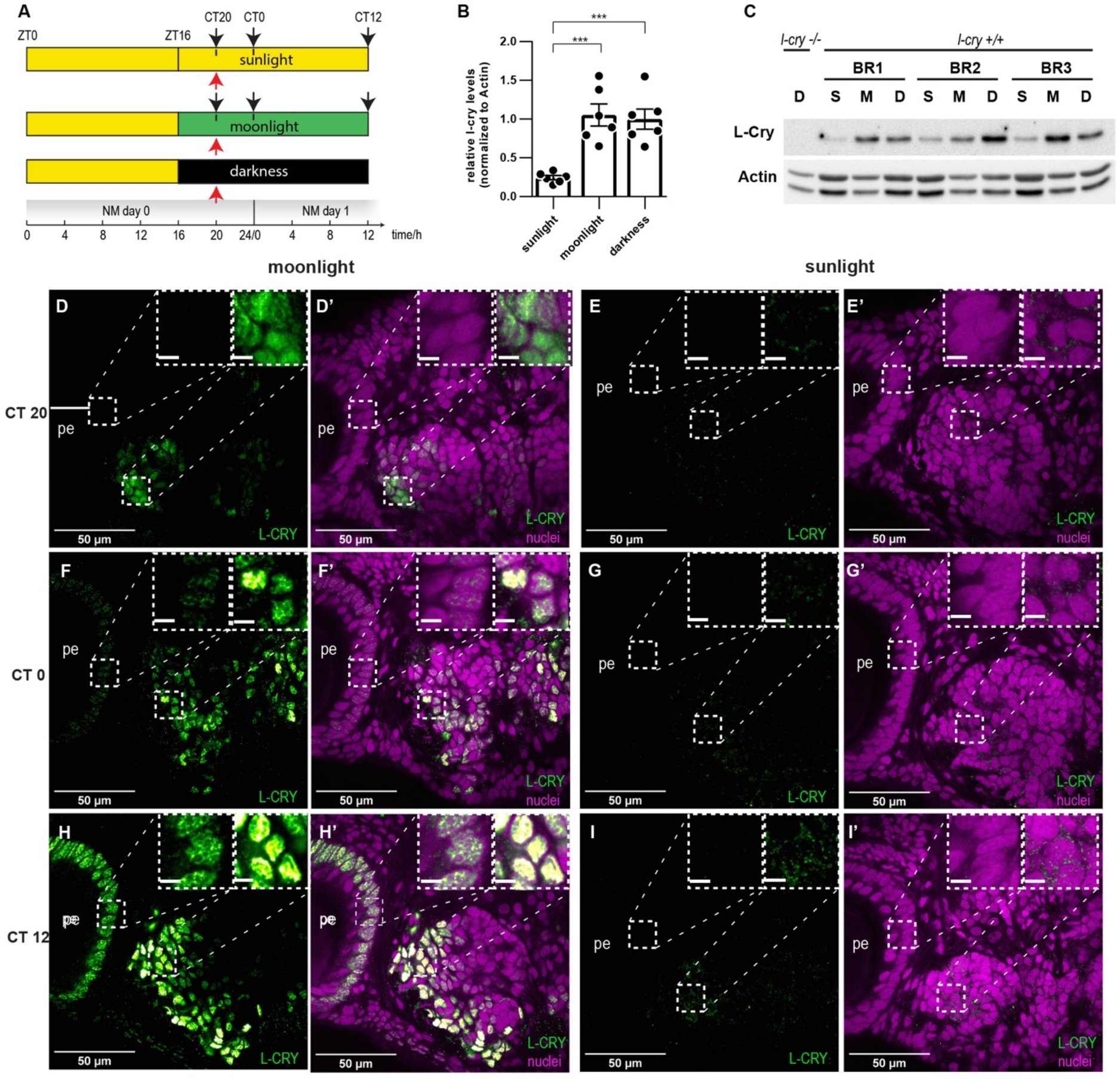
*Pdu-* L-Cry abundance and localization under darkness, naturalistic sun- and moonlight. **(A)** Sampling scheme of *Platynereis* heads for Western blot and immunohistochemistry. Red arrows: Western blots. Black arrows: immunohistochemistry. **(B)** Naturalistic sun- but not moonlight reduces L-Cry abundance. Head extracts sampled under naturalistic sunlight (S), moonlight (M) and darkness (D) were analyzed by Western blot and normalized against beta-actin, n=6 BRs. Bar graph: mean ± s.e.m. **(C)** Representative Western blot. **(D-I)** Wildtype worm heads sampled under indicated naturalistic moon- or sunlight conditions, stained with an antibody against *Pdu-*L-Cry (green). **(D’-I’)** and including nuclei stained with HOECHST (violet). Scale bar: 5μm. For comparison to dark night conditions see fig. S3 and accompanying manuscript Poehn, Krishnan et al.

Immunohistochemical analyses at two distinct time points during the first subjective night of the respective light regime (CT20; CT0, black arrows Fig. 3A), and the following mid-day point (CT12, black arrows Fig. 3A) revealed that in naturalistic moonlight, L-Cry was predominantly localized in the nuclei of the eye photoreceptors and of cells in the posterior oval-shaped brain domain (Fig. 3D–H’ and insets, for comparison to light/dark conditions: fig. S3). By contrast, residual immunoreactivity of L-Cry under naturalistic sunlight appeared to be predominantly localized to the cytosol (insets Fig. 3E–I’), in line with a sunlight-dependent degradation pathway.

These results indicate that L-Cry has the potential to signal in distinct cellular compartments to discriminate between sun and moonlight valence. This is consistent with distinct functions of L-Cry in mediating the differential impacts of sun- and moonlight on the PCC clock.

This hypothesis is further backed by biochemical data that show that naturalistic moonlight vs. sunlight results in different L-Cry photoreduction responses (see accompanying manuscript Poehn, Krishnan et al).

### Pharmaceutical disruption of canonical core circadian clock oscillations affects the PCC clock

We next wondered whether the PCC clock required the activity of the conventional core circadian clock. We previously showed that an inhibitor of the casein kinases 1δ/ε, PF670462, disrupts the worms’ core circadian clock gene oscillations (*3*). The effect of this drug on the core circadian clock has also been shown in several other aquatic animals, as diverse as cnidarian, crustacean and teleost fish species (*11*–*13*).

After validating that an incubation in 160nM of PF670462 abolished molecular oscillations of core circadian clock transcripts (fig. S4A), we assessed the effects of the drug on the timing of swarming onset. In contrast to mock-treated controls, the swarming onset in constant darkness was disrupted upon drug treatment (fig. S4B). This finding is consistent with the notion that at least a subset of canonical circadian clock genes is required for the PCC clock, although we can at present not rule out that this effect could be caused by other targets of casein kinases 1δ/ε.

### dCry prevents the fly’s circadian clock from misinterpreting moonlight

As a regular nocturnal stimulus, moonlight reaches aquatic and terrestrial habitats. The ability to properly discriminate between moon– and sunlight is therefore likely important for any species that uses light-sensitive clocks. In many species, the conventional circadian clock should likely run with a constant period, irrespective of lunar phase. Thus, moonlight would need to be “blocked” from interfering with circadian rhythmicity in those organisms. Indeed, whereas fruit fly circadian behaviour can be experimentally entrained to LD cycles with light below full moon light intensity (*14*, *15*), and constant light at moonlight intensity can extend the period length of wildtype flies (*16*, *17*), moonlight does not cause major effects on the circadian clock when combined with a LD cycle in this species (*18*–*21*).

Given our results about the importance of *Platynereis* L-Cry in discriminating between naturalistic sun-versus moonlight, and *Drosophila* dCry being its direct 1:1 ortholog, we hypothesized that this in principle functionality of the d/L-Cry family might also be present in *Drosophila melanogaster*. Specifically, we wondered if nocturnal light mimicking moonlight would cause an increased shift of the circadian clock in *dCry* mutant flies compared to controls.

We monitored locomotor behaviour of both “cantonized” *cry^01^* (*22*) and CantonS wildtype flies under LM conditions, adapting an existing locomotor paradigm (*23*), and using an artificial moonlight source matching full moon light intensities measured on land (fig. S2D,E). In wildtype flies, moonlight delayed the evening peak to 2.2h± 0.13h (mean ± s.e.m.) after night onset (Fig. 4A,C), in line with previous observations (*19*), while *cry^01^* mutants exhibited a significantly stronger delay, with the evening activity peak shifting to 4.4h ± 0.11h (mean ± s.e.m.) after night onset (Fig. 4B, C).

**Figure 4.**
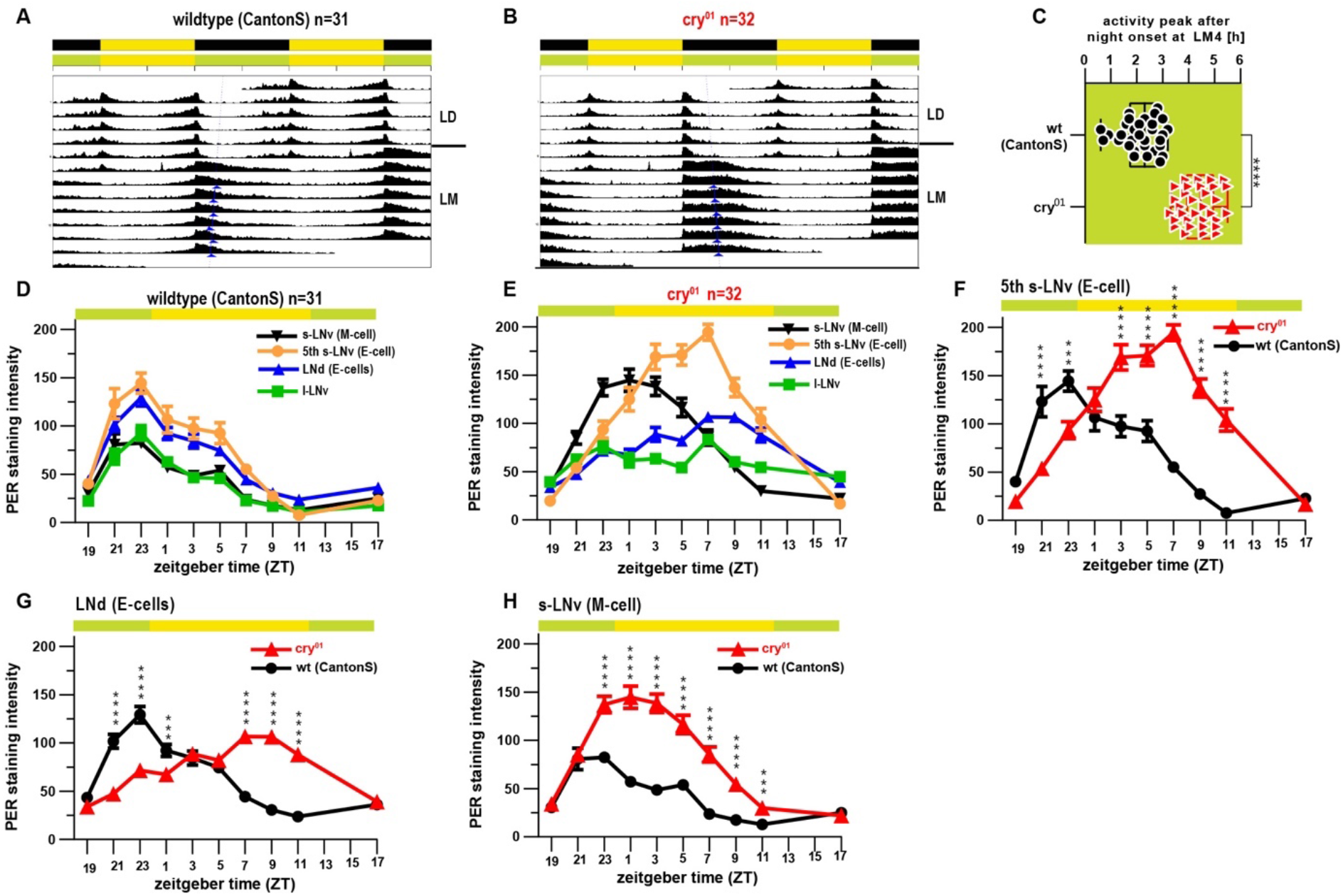
Drosophila cry protects circadian oscillator synchrony against moonlight. **(A,B)** Double-plotted actograms depicting average activity of wildtype (A) and *cry^01^* (B) flies subjected to 12:12h light:dark (LD) cycles followed by light:moonlight (LM) cycles. Blue arrowheads indicate acrophases of the respective activity rhythms. **(C)** Timing of the E-peak during LM4, calculated from the data shown in (A) and (B). The value 0 represents the time of lights off **(D,E)** Quantified anti-PER immunolabeling intensity in different groups of lateral circadian clock neurons under LM conditions (LM4) in wild-type (c) and *cry^01^* (d) individuals. **(F,G)** Detailed comparison of PER oscillations for neurons controlling evening activity, reveal a pronounced phase delay of about ~8h in cry^01^ mutants; **(H)** while neurons controlling morning activity show a more modest phase delay (2h-4h). *** : p<0.001; **** : p<0.001 ANOVA followed by Sidak’s multiple comparison test.

The increased delay of the evening activity peak in *cry^01^* mutants could either be caused by acute effects of artificial moonlight on behaviour or by a shift in the fly’s circadian clock. In order to discriminate between these possibilities, we subjected flies to artificial LM conditions and used an established immunolabeling strategy to systematically assess, over 10 distinct time points, changes in the abundance of the core circadian clock protein Period (PER) in the lateral neurons harboring the fly’s circadian pacemaker. Anatomical location and the presence or absence of immunoreactivity against the neuropeptide PDF allowed us to quantify Period abundance in l-LN_v_s, s-LN_v_s (below also referred to as morning/M-cells), as well as 5^th^ s-LN_v_s and LN_d_s (clusters harboring the evening/E-cells) (Fig. 4D-H).

Quantification across 132 CantonS wildtype individuals exposed to LM conditions revealed that oscillations of Period protein levels in the different sub-clusters were in synchrony with each other (Fig. 4D). In contrast, the corresponding *cry^01^* mutants exhibited pronounced desynchronization of Period protein oscillations between cell groups, with E-cells differing from M-cells by ~ 6h (Fig. 4E). Similar analyses of *cry^01^*-mutant flies raised in various LD cycles have not revealed such desynchronization (*24*), indicating that the effects we observed were specifically caused by exposure to artificial moonlight. When comparing Period protein abundances for the different cell classes between *cry^01^* mutants and wildtypes, Period levels in E-cells exhibited a stronger peak delay (~8h; Fig. 4F,G) than M-cells (~2h; Fig. 4H). This correlates with the fact that the peak of evening activity is significantly delayed in our behavioural analyses of cry^*01*^ mutants compared to wildtypes under LM (Fig. 4A,B). Taken together, these results indicate that the increased delay of the evening activity peak in cry^*01*^ mutants under a LM light regime is the result of a desynchronization of the circadian clock rather than an acute light effect. This suggests that *Drosophila* dCry is naturally required to reduce the effects of moonlight on circadian clock oscillations, in particular in the cell clusters harboring the evening oscillator.

### L-Cry, but not dCry is highly sensitive to moonlight

Given the genetic requirement of both L-Cry and dCry to correctly interpret moonlight under a combined moonlight/sunlight regime, we next wondered if the biochemical light sensitivity of both orthologs was also comparable. For this we purified both proteins in the presence of their co-factor flavine adenine dinucleotide (FAD) and tested for changes in absorbance after illumination. When light is sensed by dCry (*25*) or L-Cry (accompanying paper), it changes the oxidized FAD to the reduced anionic radical FAD°^-^ form, visible in the proteins’ absorbance spectrum (*25*). Extending work of the accompanying manuscript, we find that *Platynereis* L-Cry does not only respond to naturalistic full moon light (see accompanying manuscript Poehn, Krishnan et al), but does this even at intensities corresponding to 30% of full moon intensity at 4-5m seawater depths (Fig. 5A).

**Figure 5.**
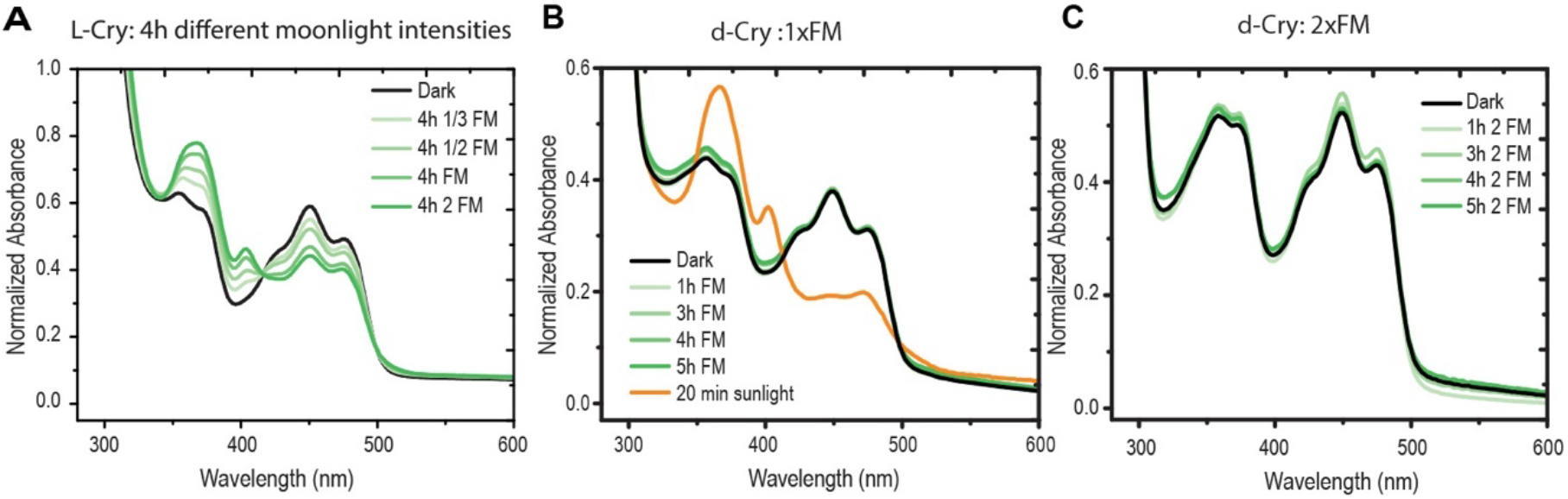
Comparison of L-Cry and dCry light detection. Illumination of purified L-Cry protein with different moonlight intensities (green) for 4h results in photoreduction (FAD°^-^ formation). FM= full moon: naturalistic full moon intensity (9.7×10^10^ photons/cm^2^/s), 1/3 FM: one third, ½ FM: one half, 2 FM: double of FM intensity. **(B,C)** dCry stimulation by moonlight (green) with naturalistic FM intensity (B) or double FM intensity (C) does not result in photoreduction, while naturalistic sunlight (yellow) does. For detailed analyses on Pdu-L-Cry responses to naturalistic sun and moonlight see accompanying manuscript Poehn, Krishnan et al.

In contrast, dCry completely failed to respond to naturalistic moonlight levels equivalent to – and exceeding – those eliciting responses in *Platynereis* L-Cry (compare Fig. 5A with B,C). However, dCry was activated by naturalistic sunlight, reaching complete FAD reduction within 20min (Fig. 5B) as observed for L-Cry (see accompanying paper), underscoring the integrity of the purified dCry protein and the functionality of the assay.

Even though dCry’s sensitivity to dim light might be higher in its cellular context (*26*), this result clearly points at differences in the molecular mechanisms between dCry and L-Cry functions. This might be well connected to the different meanings that moonlight has as an environmental cue for the daily behavior of flies versus swarming worms: Whereas fly circadian biology is likely optimized to buffer against the effect of moonlight, *Platynereis* worms, as shown above, use moonlight to precisely adjust their nocturnal swarming time to a favorable dark time window.

### R-opsin1 detects moonrise to optimize the time of swarming onset

The retention of moonlight sensitivity in *Platynereis l-cry* mutants (as evidenced by the different mutant responses under the combined moon-and sunlight regimes versus no-moonlight regimes, Fig 2D-G) indicated the existence of one or more additional light receptors required for moonlight sensation. We reasoned that the spectral sensitivity of these photoreceptors likely includes the blue-green range, given the relatively high levels of blue-green light in our moonlight measurements (fig. S2A).

The gene encoding r-Opsin1 is expressed in the adult *Platynereis* eyes both during early development (*27*, *28*) and later stages (*29*). In a heterologous expression assay established for assessing photoreceptor action spectra (*30*), *Platynereis* r-Opsin1 exhibits an irradiance response peak in the blue range (λ_max_= app. 470nm) (*31*), similar to the peak of its human melanopsin homolog. When we assessed the respective sensitivities of both receptors in side-by-side comparisons, the half-maximal effective irradiation (EI_50_) of *Platynereis* r-Opsin1 (2,3×10^10^ photons cm^−2^s^−1^) was ~100 times lower than that of melanopsin (2,5×10^12^ photons cm^−2^ s^−1^; Fig 6A), indicating a remarkably high sensitivity of *Pdu-* r-Opsin1.

**Figure 6.**
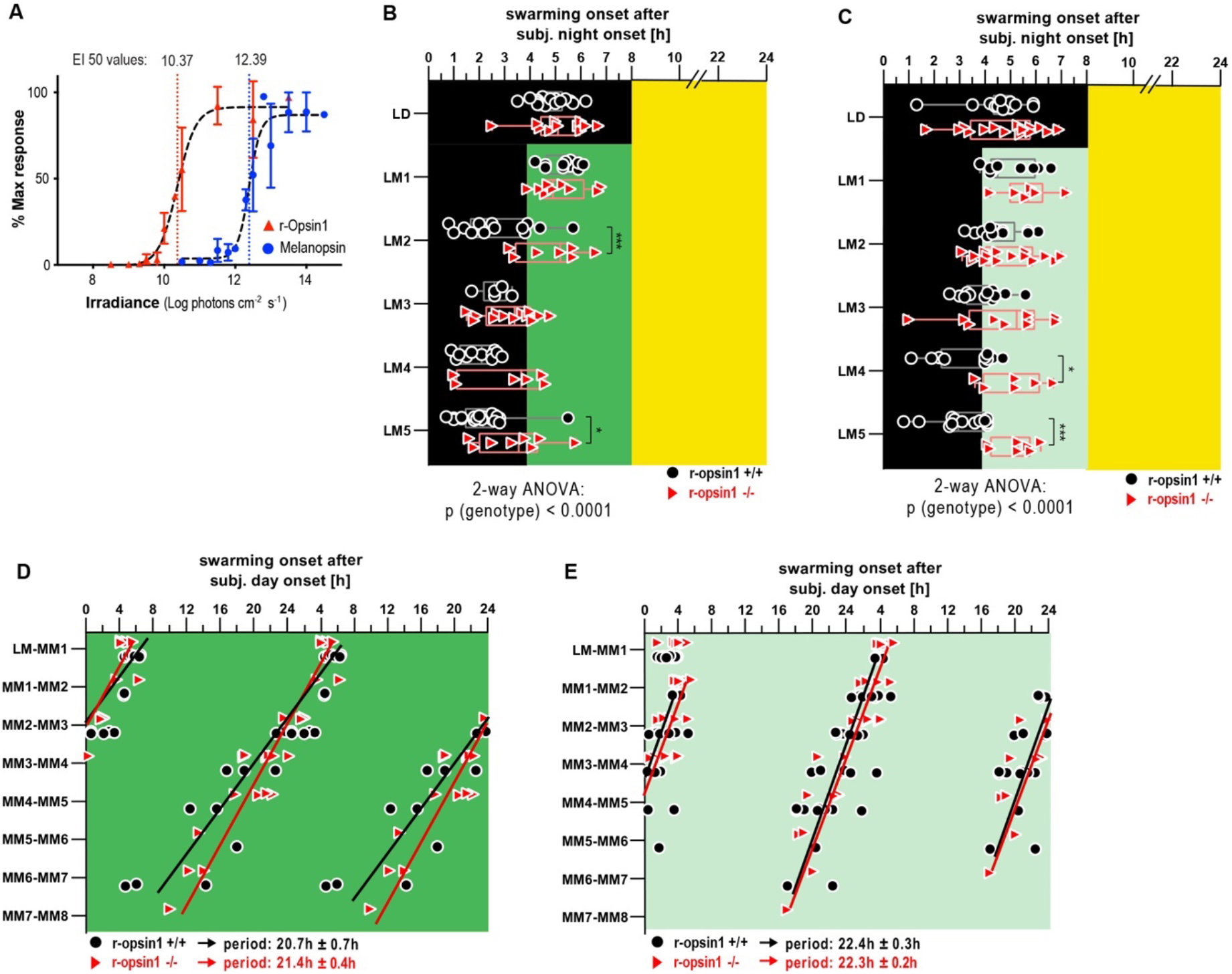
*Pdu*-r-Opsin1 functions as highly light-sensitive photoreceptor to adjust swarming onset under a waning moon light timing. **(A)** Responses of *Pdu-r-Opsin1* (red) and human Melanopsin (blue) to different blue light intensities (480nm ± 10nm) as quantified by a cell-based bioluminescent assay, reveal an ~100-fold higher sensitivity of Pdu-r-Opsin1. **(B-E)** Swarming onset of *r-opsin1−/−* and *r-opsin1+/+* worms entrained to 16:8h LD cycles and then subjected to a moon light regime typical for waning moon, i.e. moonlight during the second half of the night (LM) **(B,C)** or to constant moonlight **(D,E)** either with full moon light intensity (dark green) **(B,D)** or waning moon light intensity (light green = 20 % of full moon light intensity) **(C,E)**. * : p<0.05; ** : p<0.001; *** : p<0.0001 2-way ANOVA followed by Sidak’s multiple comparison test. Black and red lines in (D,E) indicate linear regression lines of wildtype and *r-opsin1−/−* mutants, respectively. The period length was calculated based on the slope of the regression line (from MM1-MM8) ± the 95% CI of the slope.

In the animal, this molecular sensitivity is combined with a high abundance of r-Opsin1: On the transcript level, a cellular profiling analysis revealed that *r-opsin1* is one of the topmost expressed genes in *Platynereis* adult eye photoreceptors, outnumbering a distinct co-expressed opsin – *r-opsin3* – by nearly three orders of magnitude (*31*). Moreover, in the course of the metamorphic changes that occurs during the days immediately prior to swarming, the outer segments of the eye photoreceptors – where Opsin molecules are concentrated in tightly packed membrane stacks – extend to around twice their length, suggesting an even increased sensitivity (*32*). All these facts infer that r-Opsin1 acts as a particularly high-sensitive light detector at the time of swarming.

To test whether r-Opsin1 was indeed required to mediate the impact of moonlight on the timing of swarming onset, we capitalized on an existing *r-opsin1^−17/−17^* loss-of-function allele (*31*). Following the experimental design of Fig. 1E, we subjected homozygous *r-opsin1^−17/−17^* mutants and related wildtype individuals for 5 days to naturalistic moonlight during the second half of the night (Fig. 6B). *r-opsin1^−/−^* animals exhibited a significantly reduced ability to shift their swarming onset to the dark portion of the night compared to wildtypes (Fig. 6B). This difference became even stronger with naturalistic moonlight at lower intensities (as this would be the case for the natural waning moon) (Fig. 6C). Finally, we wondered if *r-opsin1* mutants would also exhibit a reduced ability to reset the PCC under constant moonlight. Under constant moonlight at naturalistic full moon (Fig.6D) or waning moon (Fig.6E) light intensities, *r-opsin1* mutants were indistinguishable from wildtype. Comparing these results with those obtained with the *l-cry^−/−^* mutants (Fig.2B) let us conclude that *r-opsin1* specifically enables the worms to detect the rise of the moon to align the PCC accordingly.

Taken together, our data argue for two distinct roles of L-Cry and r-Opsin1 in decoding naturalistic moonlight and adjusting the PCC (Fig. 7): L-Cry, with its biochemically distinct “moonlight-state”, yet slow activation kinetics *in vitro* (see Poehn, Krishnan et al), is able to shorten the period of the PCC under sustained moonlight conditions, as they occur around natural “full moon” phases (Fig.7A). In turn, r-Opsin sensitivity, response kinetics and abundance in the eye photoreceptors make it suited to detect even weak, acute dim light, as caused by the rising moon in a “waning moon” phase, and advance the PCC (Fig. 7B). We hypothesize that the distinct nuclear localization of L-Cry in eye photoreceptors even in moonlit nights (Fig. 3) provides the necessary distinction (night/day) for activated eye photoreceptos to decode the specific valence of such nocturnal light stimuli (Fig. 7).

**Figure 7.**
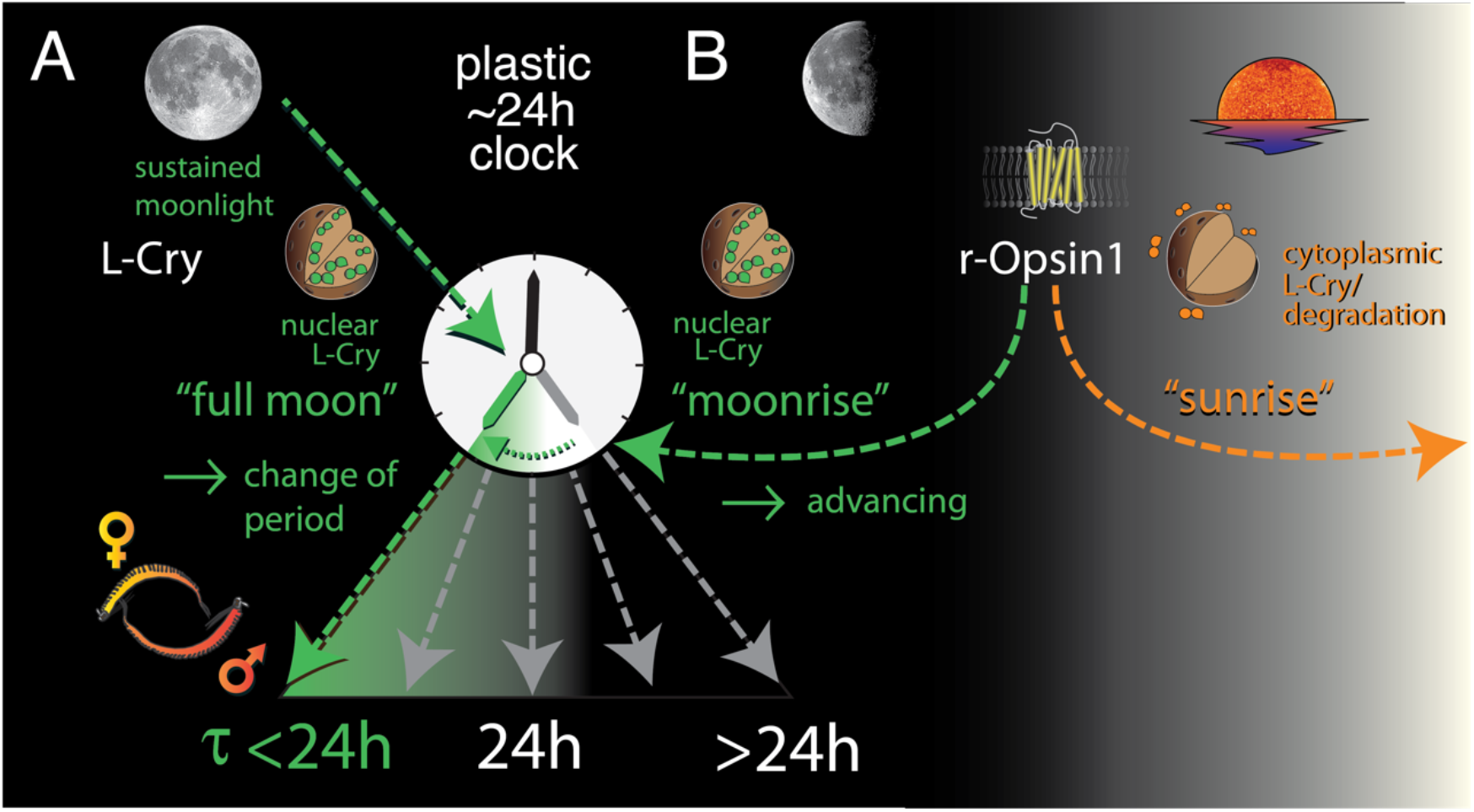
Schematic model of how the combinatorial responses of L-Cry and r-Opsin1 to naturalistic moonlight can adjust the PCC to schedule reproductive behaviour to dark portions of the night. **(A)** Sensation of sustained moonlight (“full moon”) requires L-Cryprochrome (L-Cry) and shortens the period length (τ) of the PCC. **(B)** r-Opsin1 is required to sense acute dim nocturnal light, as generated by the moonrise during the waning moon phase, and advances the PCC; correct interpretation of such acute dim nocturnal light (i.e. “moonrise” vs. “sunrise”) requires L-Cry, a function likely tied to the distinct subcellular localisation of L-Cry during night and day. Functions of sunlight in adjusting the PCC are not part of this scheme.

## Discussion

Here we uncover a ~24hr endogenous oscillator in marine broadcast-spawning worms that exhibits marked, moonlight-dependent plasticity in its period length. Its modulation by naturalistic moonlight provides a plausible model for how worms synchronize their nuptial dance, targeting a specific hour during the dark portion of moonlit nights. Restricting swarming behaviour to the dark portion of the night might be advantageous to avoid predators that hunt during moonlight. On a mechanistic level we suggest that this PCC clock shares elements with the conventional core circadian oscillator and reveal two highly sensitive light receptors, r-Opsin1 and L-Cry, that are critical to sense and interpret naturalistic moonlight.

Sensitivity to moonlight is directly relevant for a broad panel of marine broadcast spawners. The challenge of “tagging” nocturnal light information with the correct valence, however, likely extends beyond this specific ecological context. The classical categorization of organisms into nocturnal versus diurnal species (*33*, *34*) typically neglects the aspect of moonlight. Any animal entraining its ~24hr clock to light will need to correctly interpret the occurrence of nocturnal light. Even though it has been shown that the circadian system of many species is sensitive to light levels as low as moonlight intensity, such as in flies (*14*, *15*) and mice (*35*), chronobiological studies have so far put relatively little effort in dissecting how animal clocks prevent potential disturbance of moonlight, and interpret naturalistic light regimes that combine both sun– and moonlight.

The data presented in this and the accompanying manuscript provide possible mechanistic explanations for the ability of the PCC clock to decode a combined sun and moon light regime. A first tier is connected to the specific properties of cryptochrome: While under naturalistic moonlight, *Platynereis* L-Cry protein levels remain elevated, comparable to dark conditions, and are predominantly localized to the nucleus, the onset of sunlight causes a rapid degradation, with residual L-Cry protein found in the cytoplasm. On the biochemical level, L-Cry is highly sensitive to naturalistic moonlight. Moonlight evokes a different state in L-Cry than sunlight (see extensive comparison of sunlight vs. moonlight in Poehn, Krishnan et al). Taken together, these data are consistent with the idea that – besides the canonical strong-light induced degradation-based signaling pathway for cytoplasmic Cryptochrome – L-Cry possesses a second, dim-light induced, nuclear mode of signaling. A second lead is provided by our identification of r-Opsin1 as a second moonlight sensor. It remains to be uncovered, however, how the r-Opsin1-dependent signals tie in with the different signaling states of L-Cry.

Evidence for plasticity of the conventional circadian clock has started to emerge from other marine systems: Work on the circatidal oscillators of oysters maintained under controlled lab conditions revealed that core circadian clock genes exhibit ~12.4hr cycles under constant darkness, while the transcripts of the same genes cycle with a ~24hr oscillation under light/dark conditions (*36*). This provides evidence for the ability of the canonical clock to alternate between circadian (~24h) and (semi)circalunidian (~12.4h/~24.8h) periodicities. Interestingly, switches between circadian and circalunidian cycles might also occur in humans. It was shown that mood switches of bipolar patients correlate with a period lengthening of their body temperature cycles that looks as if the circadian timing system can be intermittently entrained to a 24.8h rhythm (*37*). While such observations in human remain highly enigmatic, we anticipate that research on organisms for which lunar impact is of known biological relevance will be key to disentangle the interplay of solar and lunar timing cues.

## Material and Methods

### Worm culture

Worms were grown as described previously (*38*). In short: worms were kept in plastic boxes filled with a 1:1 mixture of natural sea water and artificial sea water (30% Tropic Marine) and exposed to a 16h : 8h light:dark light regime. To entrain their circalunar clock, worms receive 8 nights of continuous nocturnal light each month to mimic full moon (FM).

Strains: *l-cry^−/−^:* homozygous Δ34, generated in the VIO-strain background (see accompanying manuscript Poehn, Krishnan et al.). Wildtype worms used for comparison to *l-cry^−/−^* worms are cousin relatives to *l-cry^−/−^* worms.

*r-opsin1^−/−^:* homozygous Δ17, generated in the *r-ops1::GFP* transgenic strain (*31*). Wildtype worms used for comparison are from the *r-ops1::GFP* transgenic strain from which the mutant was generated.

### Natural light measurements

Under water measurements of natural sun- and moonlight at the habitat of *Platynereis* were acquired using a RAMSES-ACC-VIS hyperspectral radiometer (TriOS GmbH) for UV to IR spectral range (see (*7*) for details). Radiometers were placed at 4m and 5m water depth close to *Posidonia oceanica* meadows, which are a natural habitat for *P. dumerilii*. Measurements were recorded automatically every 15min across several weeks in the winter 2011/2012 (at 5m depth) and during spring 2011 (at a 4m depth). To obtain an exemplary sunlight spectrum, the sunlight measurements taken at 5m depth between 10 am-4 pm on 25.11.2011 we averaged. To obtain a full moon spectrum for the 5m depth location measurements taken from 10pm to 1am on a clear full moon night (10-11.11.2011) were averaged. To control for technical noise caused by the measurement device at these low light intensities, a NM spectrum was obtained by averaging measurements between 7:15pm to 5am on a NM night on 24.11.2011, and subtracted from the FM spectrum. The resulting spectrum is plotted in fig S2A. To validate that this spectrum is representative of a typical full moon spectrum at the habitat of *Platynereis*, we averaged moonlight measured between 10:15 pm to 2am during a full moon night (17.-18.04.2012) and subtracted a NM spectrum measured two weeks earlier from 4m depth (fig. S2A). To benchmark these moonlight spectra measured under water with moonlight measured on land, we compared the underwater spectra to a publicly available full moon spectrum measured on land on 14.04.2014 in the Netherlands (fig.S2A, http://www.olino.org/blog/us/articles/2015/10/05/spectrum-of-moon-light). As expected, light with longer wavelengths was strongly reduced in the underwater measurements compared to the surface spectrum, since light with longer wavelengths penetrates water less efficiently.

### Behavioural setup and analyses of swarming onset

All behavioural experiments, except Fig. 1B and Fig. 2F,G were performed with worms that received LD conditions without any nocturnal light (FM) for at least 9 days. Since most *l-cry* mutants spawn during the first 9 nights after the FM stimulus under standard worm culture conditions (Poehn et al), the monthly FM stimulus was omitted for *l-cry* mutants and wildtypes in order to test swarming worms without confounding effect of a recent nocturnal (highly artificial) light stimulus on swarming onset. Sexually maturing worms were placed in seawater filled individual hemispherical concave wells (diameter = 35mm, depth = 15mm) of a custom-made 36-well clear plastic plate. Video recording of worm’s behavior over several days was accomplished as described previously (*3*), using an infrared (λ = 990 nm) LED array (Roschwege GmbH) illuminating the behavioral chamber and an infrared high-pass filter restricting the video camera. Worms were recorded at least until initiation of swarming (fig.S1A). Naturalistic sun- and moonlight were generated by custom made LEDs (Marine Breeding Systems, St. Gallen, Switzerland) (for spectra and intensity see fig. S2B,E). Naturalistic sun- and moonlight were used in all worm experiments, except for data obtained in Fig. 1B and Fig. 2E,F,G were we used prototype artificial sun- and moonlight LEDs (fig. S2C).

Spectra were measured with a calibrated ILT950 spectrometer (International Light Technologies Inc., Peabody, USA). To reliable measure the artificial moonlight, the detector was placed 12cm away from the moonlight source, and based on this measurement moonlight intensity was calculated using the inverse square law for worm position, which was ~51 cm away from the moonlight source.

After video recording, an automated tracking software was used to deduce locomotor activity of individual worms across the time of the recording (*7*). The exported locomotor activity trajectories, which reflect the distance moved of each worm’s center point across 6 min time bins, were analyzed in ActogramJ to manually identify the swarming onset moment. In ambiguous cases (e.g. only little movement detected) we manually analyzed the video recordings to identify the moment when a sexually mature worm left its tube, which was regarded as swarming onset. Swarming onset data were plotted and analyzed using GraphPad Prism 8.0 (La Jolla, USA). ANOVA was used to test if swarming onset was statistically different across the different days of an experiment. This was followed by Dunnetts multiple comparison test, comparing each day of the experiment with swarming onset during LD conditions. To test differences in swarming onset between mutants and wildtypes across different days of an experiment with varying light conditions, 2-way ANOVA was used followed by Sidak’s multiple comparison test. To identify the free-running periodicity under constant light conditions linear regression analysis was performed. The period length was calculated based on the slope of the regression line ± the 95% CI of the slope. Swarming onset data are presented including the individual data points and a box plot. The whiskers of the box blot represent minimal and maximal values.

### Recording of locomotor activity in *Drosophila melanogaster*

Locomotor activity was recorded under constant temperature (20°C) from 0-1 day old male Canton-S and *cry^01^* (CantonS background) flies using the *Drosophila* Activity Monitors from Trikinetics Incorporation (Waltham, MA, USA)(*23*). Flies were first recorded for 5 days under 12h light - 12h dark cycles (=LD with ~100 l× standard white light LED), and then under for 7 days under 12h light – 12h artificial moonlight cycles (=LM cycles; for spectrum and intensity of artificial moonlight see fig.S2C). The average actograms and the centers of maximal activity were calculated and plotted with ActogramJ(*39*). The phases of evening activity maxima under LM conditions were determined using the ActogramJ tool “acrophase”. To test for differences in the acrophase of wildtype and *cry^01^* flies at LM4, an unpaired student-test was performed.

### Western blots

Four anaesthetized worms were decapitated and heads transferred to a 1.5ml tube containing 150 μl RIPA lysis buffer (R0278 Sigma-Aldrich) supplemented with 10% Triton X100 and protease inhibitor (cOmplete Tablets, EDTA-free, EASYpack, Roche) per biological replicate. The tissue was homogenized by grinding using a tightly fitting pestle. All steps on ice. Cell debris was pelleted by centrifugation. Protein concentration of lysates was determined using Bradford reagent (BIORAD). Proteins were separated by SDS-gel electrophoresis (10% Acrylamide) and transferred to nitrocellulose membrane (Amersham™ Protran™ 0,45μm NC, GE Healthcare Lifescience). Quality of transfer was confirmed by staining with Ponceau-S solution (Sigma Aldrich). After 1h of blocking with 5% slim milk powder (Fixmilch Instant, MARESI) in 1xPTW (1xPBS/0.1% TWEEN 20) at room temperature, the membrane was incubated with the appropriate primary antibody, diluted in 2.5% milk/PTW at 4°C O/N. [anti-L-Cry 5E3-3E6-E8 (1:100) and anti-L-Cry 4D4-3E12-E7 (1:100); anti-beta-Actin (Sigma, A-2066, 1: 20.000)]. After 3 rinses with 1xPTW the membrane was incubated with the species specific secondary antibody [anti-Mouse IgG-Peroxidase antibody, (Sigma, A4416, 1:7500); Anti-rabbit IgG-HRP-linked antibody (Cell Signaling Technology, #7074, 1:7.500] diluted in 1xPTW/1% slim milk powder for 1 hour. After washing, SuperSignal™ West Femto Maximum Sensitivity Substrate kit (Thermo Fisher Scientific) was used for HRP-signal detection and finally signals were visualized by ChemiDoc Imaging System (BIORAD). Bands were quantified in “Image Lab 6.1” (BIORAD)

### Immunohistochemistry

Portions of *Platynereis dumerilii* bodies containing head and jaw were dissected and fixed in 4% PFA at 4° C for 24 h. Afterwards, methanol washes at room temperature (r.t., shaking) and a 5-minutes long digestion using Proteinase K (r.t., not shaking) were employed as means of permeabilization. The worm heads and jaws were then post-fixed with 4% PFA for 20 min at r.t. and washed using 1× PTW (PBS-0.1% Tween 20^®^ (Sigma Aldrich)) 5 times for 5 min. This was followed by over-night incubation in a hybridization mixture(*40*), commonly used for in situ hybridization (at 65° C in water bath; the solution exchanged once, after the first hour of incubation). Several washing steps were performed the following day, at 65° C in a thermo-block, not shaking (washing sequence, solutions and durations: a. 2 times 20 min with 50% formamide/2X standard saline citrate - 0.1% Tween 20^®^ (Sigma Aldrich), SSCT; b. 2 times 10 min with 2X SSCT; c. 2 times 20 min with 0.2X SSCT). Samples were subsequently blocked using 5% sheep serum (Sigma-Aldrich) (r.t., 90 min, shaking) and incubated for at least 36 h (4° C, shaking) in a mixture of two monoclonal antibodies against L-Cry, 5E3-3E6-E8 and 4D4-3E12-E7 (1:100 and 1:50, correspondingly, in 5% sheep serum (Sigma-Aldrich)) (see accompanying manuscript for further details). Next, samples were washed with 1× PTW 3 times for 15 min (r.t., shaking) and a 1 time over night (4° C, shaking). A Cy3 goat anti-mouse IgG secondary antibody (A10521, Thermo Fisher Scientific) was added in dilution 1:400 in 2.5% sheep serum to specifically detect the bound primary antibody (incubation time and conditions, as well as the following washing steps, were the same as those of the primary antibody). To label nuclei, samples were incubated for 30 min in Höchst 33342 (H3570, Thermo Fisher Scientific), diluted 1:2000 (r.t., shaking), washed 3 times for 15 min using 1× PTW and mounted in 87% glycerol (Sigma-Aldrich)/ddH_2_O containing 25 mg/ml DABCO (Roth/Lactan). All solutions were made using 1× PTW unless stated otherwise.

Imaging of the worm heads was done using a Zeiss LSM 700 laser scanning confocal microscope and LD LCI Plan-Apochromat 25X and Plan-Apochromat 40X by CHD objectives, T-PMT detection system and Zeiss ZEN 2012 software (lasers used: DAPI 405 nm and Cy3 555 nm). Image analysis was performed using the software Fiji/ImageJ (*41*).

### Period oscillations in Drosophila clock neurons

To compare the effect of moonlight between cry mutants and wildtypes on the Period oscillations in the different clock neuron clusters we entrained 0-1 day old male Canton-S and *cry01* (CantonS background) flies first under 12h light - 12h dark cycles (~100 l× standard white light LED), and then subjected them to artificial moonlight during the night (=LM cycles; for spectrum fig.S2C) for another 4 days. At LM4 whole flies were fixed at the indicated ZTs (for 3h) with 4% PFA + 0.1% TritonX100. Flies were then washed 3×10min in PBT 0.5% and their brains were dissected. Subsequently, brains were blocked with 5% NGS in PBT 0.5% for 3 hours. Brains were incubated for 48h at 4°C with the following primary antibodies diluted in PBT 0.5% + 5% NGS: rabbit anti-PER (1:1000), mouse anti-Pdf (1:1000). The secondary antibodies were goat anti-rabbit Alexa^TM^ fluor 488 (1:200) and goat anti-mouse Alexa^™^ 635 (1:200) incubated at 4°C overnight. Before mounting, brains were washed 6x with PBT 0.5% (last wash with PBT 0.1%) and then mounted in Vectashield H-1000. Images were acquired with TCS SPE Leica confocal microscope using a 20-fold glycerol immersion objective (Leica Mikrosystems, Wetzlar, Germany) and analyzed with ImageJ as described in ref. (*42*). PER staining intensity in the different pacemaker cell groups was examined in 12-15 brains (one hemisphere per brain) per timepoint and genotype. To obtain PER staining intensity above background for of each cell group, the PER signal of all cells of a cell group in one hemisphere was averaged and background signal measured near this cell group was subtracted. In case not all cells of a specific cell group could be identified, these missing cells were ignored for analysis.

Finally, to obtain an average staining intensity per cell group, the corresponding staining intensities of all 12-15 brain hemispheres sampled during one timepoints were averaged.

### Opsin spectral sensitivity comparison

To investigate the spectral sensitivity comparison of Pdu r-opsin1 to human melanopsin, mammalian expression vectors for both opsins were independently co-transfected into HEK293 cells along with an expression vector containing the luminescent calcium sensitive protein, Aeuqorin (pcDNA5/FRT/TO mtAeq) using Lipofectamine 2000 to access the activation of Gαq signaling as shown in previously published work (Roger publication, Bailes et al). After 6hrs incubation, the medium was changed to DMEM containing 10% FBS and 10uM 9-cis retinal, after which point the cells were protected from light. The following day, medium was changed to L-15 without phenol red, containing 10uM Coelentrazine-h and 10uM 9-cis retinal. Individual wells were briefly exposed to a 2s flash of near monochromatic light (480nm +/− 10nm) produced from an Xenon arc lamp and delivered via a fiber-optic cable fixed ~10cm above the relevant well and accessed for increase in calcium level by measuring the raw luminescence (RLU) signal with a resolution of 0.5s and cycle of 2s. Luminescence was read using a Clariostar (BMG labtech). Light intensity was modified using combinations of 0.9, 0.2 and 0.1 Neutral density filters. RLU measured during dark incubation preceding the light pulse were used as baseline. Maximum response was determined by the peak luminescence value post light flash, normalised to the maximum luminescence value recorded, per opsin, for that experiment. The resultant maximal response value acquired from each replicate were plotted against the irradiance measured for tested wavelength. This irradiance response curve was then fitted with a sigmoidal dose response function to understand the maximum sensitivity of both opsins.

### Casein kinase inhibitor treatment and qPCRs

Worms were treated with indicated concentrations of PF-670462 for 3 days under LD conditions during new moon. For sampling, worms were first anaesthetized for ca. 10min with a 1:1 mixture of seawater and 7.5% (w/v) MgCl2 solution. The head was then cut behind the posterior eyes with a scalpel at the indicated timepoints. Five heads were pooled per biological replicate, immediately frozen in liquid nitrogen and stored at −80°C until RNA extraction.

For RNA extraction, 350μl of RNAzol RT (Sigma-Aldrich) were added to the samples and lysis was performed with TissueLyser II (Qiagen) at 30Hz for 2min. Afterwards, RNA was extracted using Directzol RNA Miniprep kit (Zymo Research) following the manufacturer’s instructions with additional on-column DNaseI digest. RNA was eluted in 34μl of nuclease-free water.

Total RNA (300ng per sample) was reverse transcribed using QuantiTect Reverse Transcription Kit (Qiagen). The resulting cDNA was diluted to a volume of 60μl. qPCR reactions were performed in 20μl total volume with Luna Universal qPCR Master Mix (New England Biolabs). Target genes and reference controls were analysed in duplicate reactions for all samples. Plate control cDNA and -RT controls were included on each plate. cdc5 was used as reference gene(*3*). Expression levels were calculated using the Δct method. Relative expression values were calculated with the formula: relative expression = 2 - Δct.

### Recombinant expression and purification of L-Cry and dCry proteins

L-Cry was expressed and purified from insect cells as described in the accompanying manuscript (Poehn/Krishnan et al). N-terminally His6-tagged dCry was expressed in *Spodoptera frugiperda* (*Sf9*) insect cells using a pFastBac HTb expression vector (Berndt et al, 2007). 1 L of 1 * 10^6^ *Sf9* cells/ml in sf900II media were transfected with P1 virus stock and incubated at 27°C for 72 h. Harvested cell pellets were resuspended in lysis buffer (25 mM Tris pH 8.0, 300 mM NaCl, 20 mM imidazole, 5% glycerol, 5 mM β-mercaptoethanol) and lysed by sonication. The lysate was centrifuged and the clarified supernatant loaded onto a 5ml HisTrap HP nickel affinity column (GE Healthcare). dCry protein was eluted with 100 mM imidazole, diluted with low salt buffer (50 mM Tris pH 8.0, 5% glycerol, 1mM DTT) and loaded onto a 5 ml DEAE sepharose anion exchange column (GE Healthcare). After gradient elution (0 to 500 mM NaCl), dCry containing fractions were concentrated and loaded onto a HiLoad S200 16/60 size exclusion chromatography (SEC) column (buffer 25 mM Tris pH 8.0, 150 mM NaCl, 5% glycerol, 1 mM TCEP). SEC fractions containing pure dCry protein were pooled, concentrated and stored at −80°C until further use. All purification steps were carried out in dark- or dim red light conditions.

### UV/VIS spectroscopy of L-Cry and dCry

UV/VIS absorption spectra of purified L-Cry and dCry proteins were recorded on a Tecan Spark 20M plate reader. An intensity calibrated naturalistic moonlight source (fig.S2C) was used for moonlight UV/VIS spectroscopy on L-Cry and dCry. Naturalistic full moon (FM) intensity was set to 9.67 × 10^10^ photons cm^−2^s^−1^. To analyze moonlight dose-dependent FAD photoreduction of L-Cry, dark-adapted L-Cry was illuminated with different moonlight intensities (1/3 FM, 1/2 FM, FM and 2 FM intensity) continuously for 4 h on ice and UV-VIS spectra (300 – 700 nm) were collected after 4 h. To analyze sunlight- and moonlight dependent FAD photoreduction of dCry, dark-adapted dCry (kept on ice) was continuously illuminated with naturalistic sunlight (1.55 × 10^15^ photons cm^−2^ s^−1^ at the sample) or naturalistic moonlight (9.67 × 10^10^ photons cm^−2^ s^−1^ at the sample) and UV-VIS spectra (300 – 700 nm) were collected at different time points.

### Statistical analyses

We used one-way ANOVA followed by Dunnett’s test to test if the timing of swarming onset during LD conditions differs compared to conditions were worms are subjected to moonlight conditions on top of a LD cycle. We used two-way ANOVA followed by Sidak’s test to test if and during which days the timing of swarming onset differs between mutant and wildtypes across different days of a behavioural experiment. To compare if two sets of data had different variances, a F-test as part of t-test statistics was performed. Swarming onset data are shown as individual data points, and additionally represented as box plots with whiskers reaching to the maximal and minimal value.

Western blot data, which assessed head L-Cry levels during sunlight, moonlight and darkness conditions were analyzed with one-way ANOVA followed by Tukey’s multiple comparison test to test for significant differences in L-Cry abundance between the different light conditions.

To compare period oscillation in the different cell groups between *cry01* mutants and wildtype flies over different ZTs we used two-way ANOVA followed by Sidak’s test.

## Acknowledgements

We thank the members of the Tessmar-Raible, Raible, Helfrich-Förster and Wolf groups for discussions. Andrej Belokurov and Margaryta Borysova for excellent worm care at the MFPL aquatic facility. We are grateful for support by the IMB Protein Production and Proteomics Core Facilities (instrument funded by DFG INST 247/766-1 FUGG).

## Funding

K.T-R. received funding for this research from the European Research Council under the European Community’s Seventh Framework Programme (FP7/2007–2013) ERC Grant Agreement 337011 and the Horizon 2020 Programme ERC Grant Agreement 819952, the research platform ‘Rhythms of Life’ of the University of Vienna, the Austrian Science Fund (FWF, http://www.fwf.ac.at/en/): SFB F78 and the HFSP (http://www.hfsp.org/) research grant (#RGY0082/2010). S.K. is a recipient of a DFG fellowship through the Excellence Initiative by the Graduate School Materials Science in Mainz (GSC 266). C.H-F. was funded by Deutsche Forschungsgemeinschaft (DFG), collaborative research center SFB 1047 “Insect timing,” Project A1 and A2. None of the funding bodies was involved in the design of the study, the collection, analysis, and interpretation of data or in writing the manuscript. RJL received support from the HFSP (project grant RGP0034/2014).

## Supplementary Figures

**Fig. S1.**
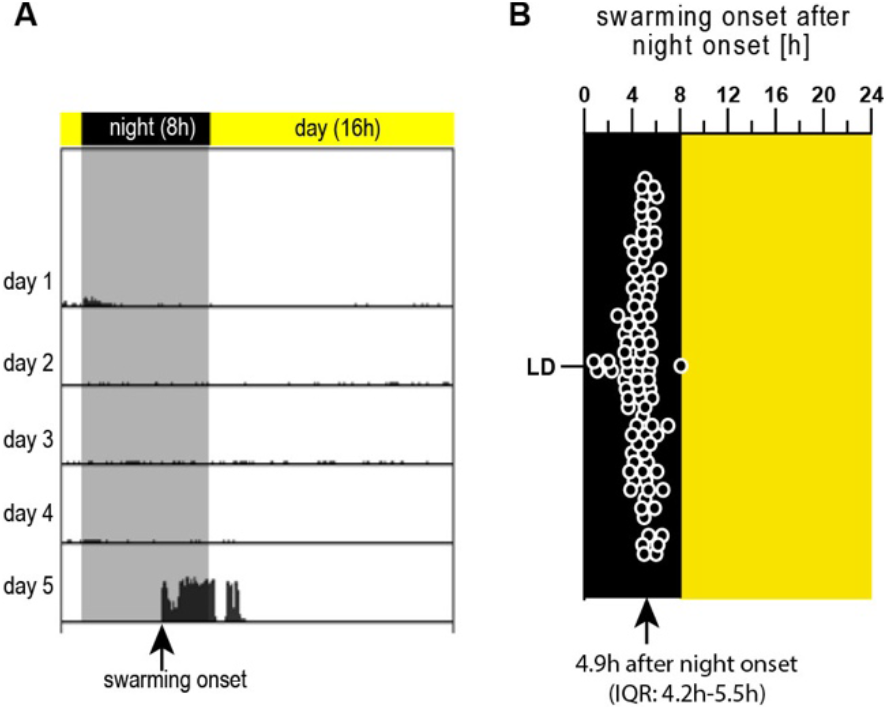
Determination of the timing of swarming onset by tracking locomotor activity. **(A)** Exemplary actogram showing locomotor activity of a sexually maturing worm during the days prior to swarming and in the night of swarming. Swarming onset is correlated with a striking increase in locomotor activity. See Supplementary Video 1. **(B)** Coordinated swarming onset of separated worms that were kept under a 16h:8h LD cycle for at least 9 days prior to swarming (n=92). Median swarming onset was 4.9h after night onset (IQR: 4.2h-5.5h)

**Fig. S2.**
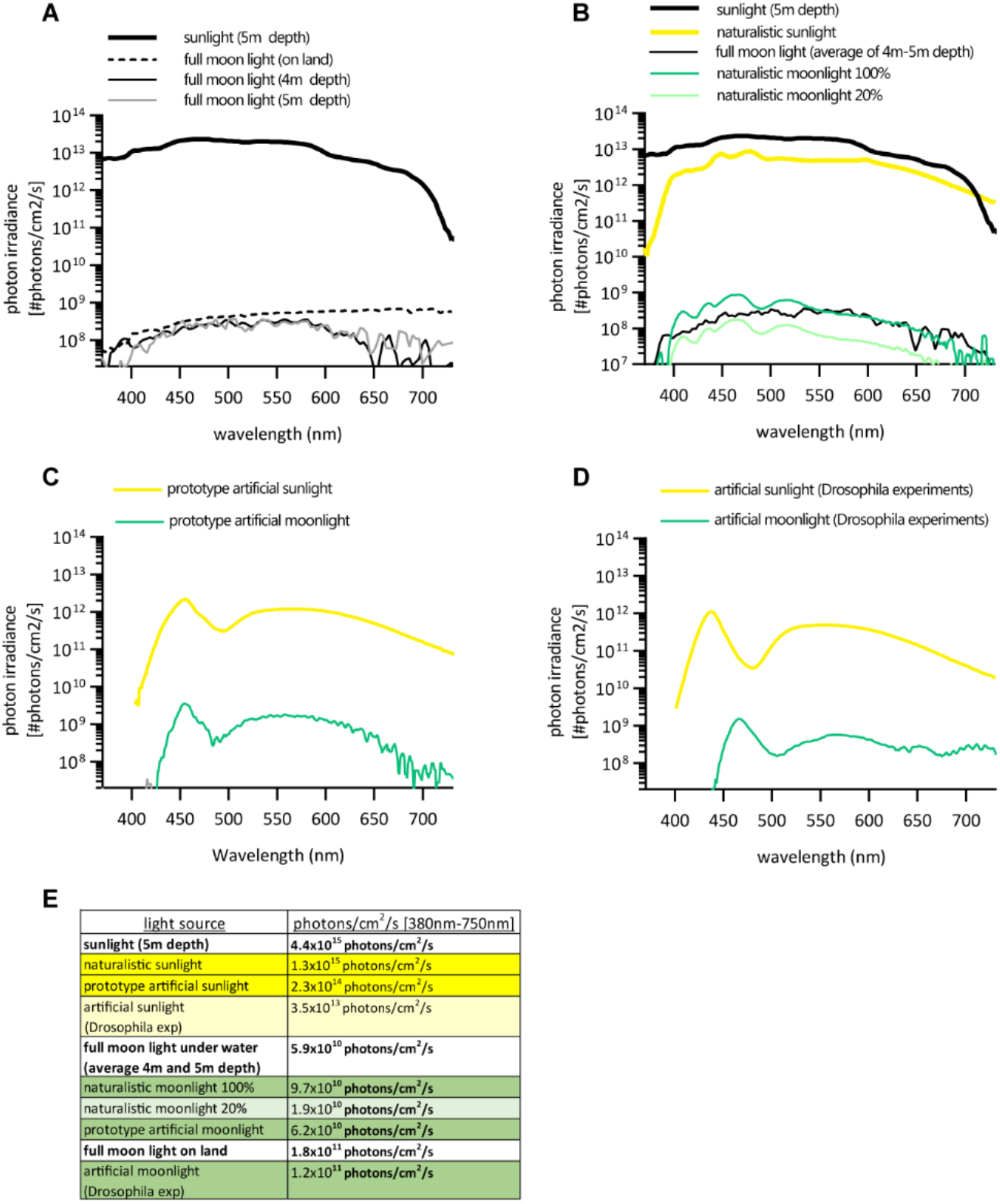
Sun- and moonlight spectra. **(A)** Exemplary natural sunlight and full moon spectra measured under water at the natural *Platynereis* habitat in the coastal waters of Ischia/Italy. Sunlight spectrum was measured at 5m water depth on 25.11.2011 (9.7×10^10^ photons/cm^2^/s [380nm-750nm)average 10am-4pm), and the two full moon spectra were measured at 4m and 5m water depth on 17.-18. April 2012 (average 10:15pm-2am) and 10.-11.2011 (average 10pm-1am), respectively. To benchmark the underwater moonlight measurements a publicly available full moon light spectrum measured on land is included (http://www.olino.org/blog/us/articles/2015/10/05/spectrum-of-moon-light). **(B)** Custom designed naturalistic sun (yellow) and moonlight spectra (dark and light green) used for all *Platynereis* experiments (except for Fig.2E, F, G and fig. S1) compared to natural sun and moonlight spectra. **(C)** Prototype artificial sun- and moonight spectra used for experiments shown in Fig. 1B and Fig.2 E,F,G. **(D)** Artificial sun and moonlight experiments used for Drosophila experiments. **(E)** Total light intensities of the spectra shown in (A-D). All spectra reflect light intensities at the distance relevant for experiments.

**Fig. S3.**
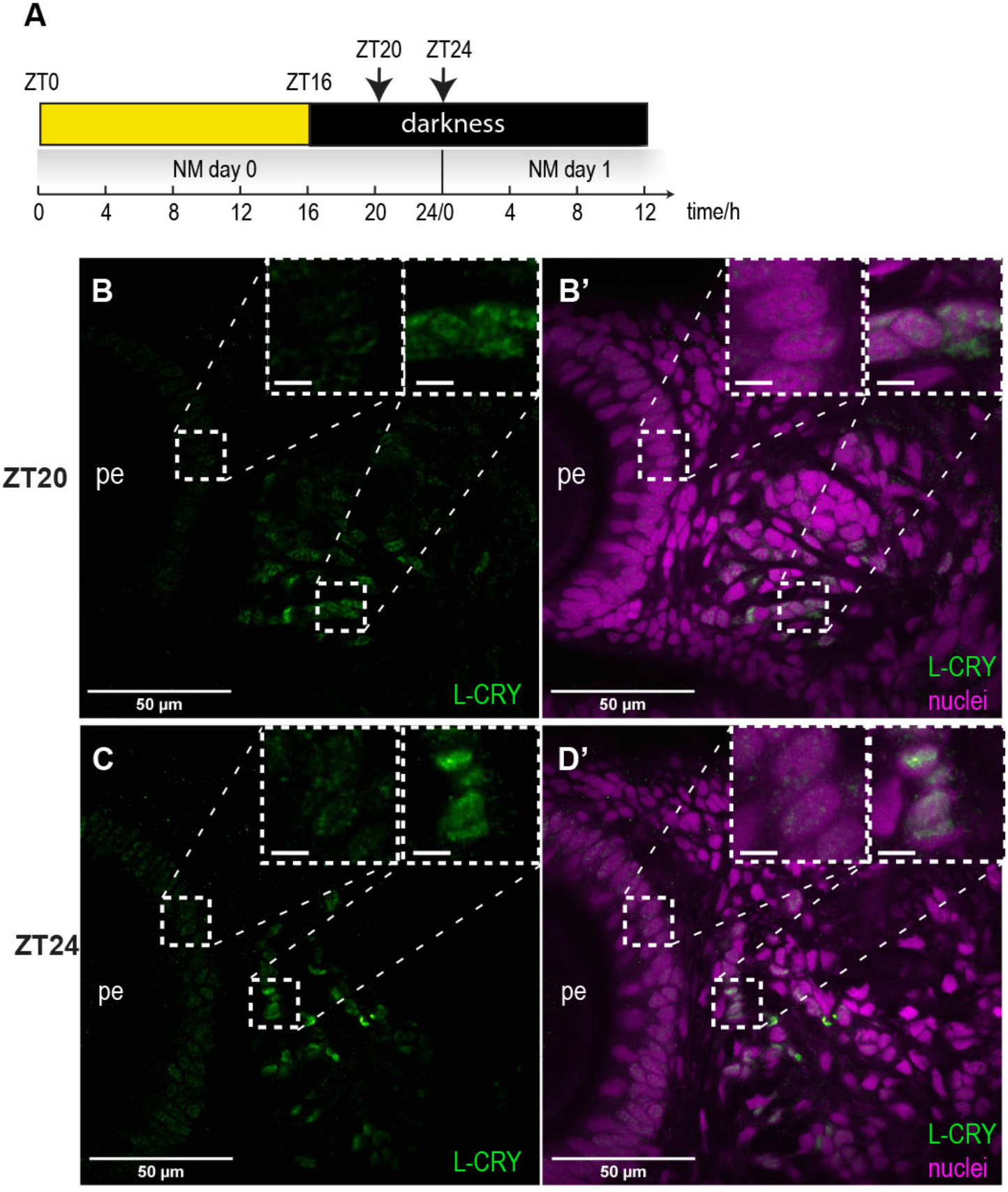
L-Cry localizes to the nucleus during dark nights. **(A)** Sampling scheme of *Platynereis* heads for immunohistochemistry. **(B,C)** *Pdu-*L-Cry (green); **(B’C’)** *Pdu*-L-Cry including nuclei stained with HOECHST (violet). For further details see Fig.3.

**Fig. S4.**
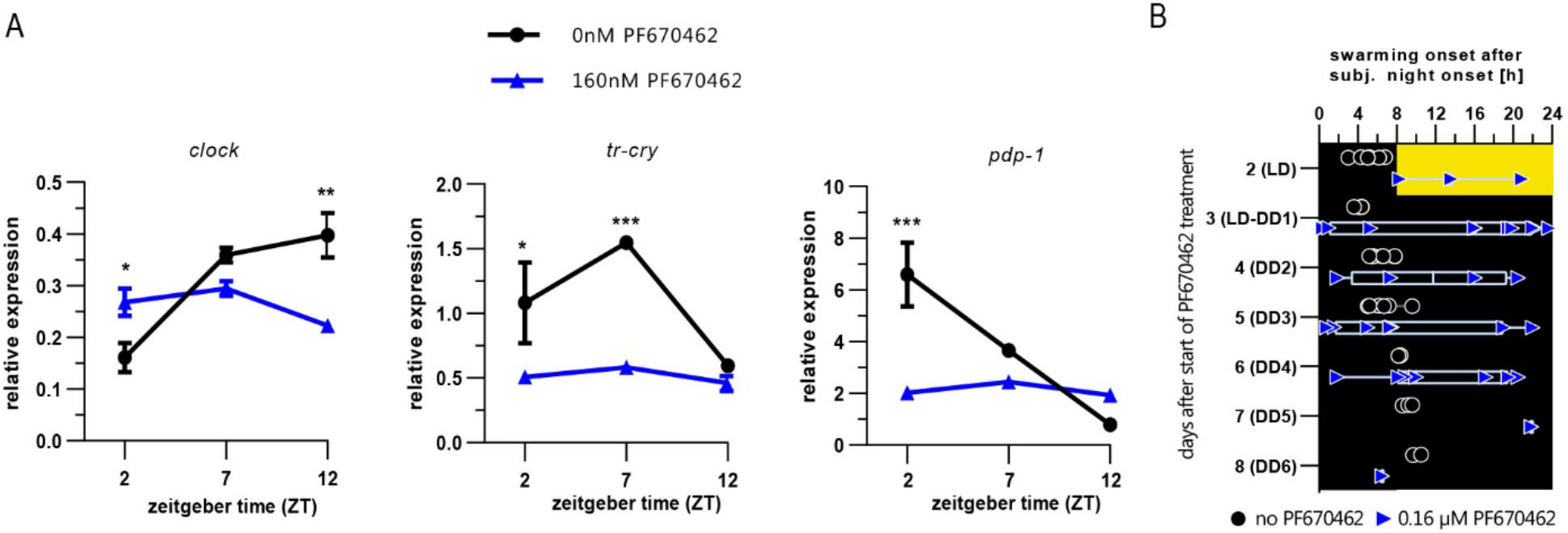
Treatment with a casein kinase 1δ/ε inhibitor disrupts circadian clock oscillations and synchronized swarming onset. **(A)** Treatment of 160nM of casein kinase 1δ/ε inhibitor PF670462 results in severely disrupted circadian clock gene transcriptional oscillations in head extracts of premature worms. Expression levels are normalized to *cdc5* levels. **(B)** Swarming onset of worms after at least 9 days after last FM stimulus under LD followed by DD conditions treated with the casein kinase 1δ/ε inhibitor PF670462 (blue triangles); untreated references (black dots) include individuals also shown in Fig. 1C. Values are means ± SEM; n = 3BRs with 4-5 heads/BR. * : p<0.05; ** : p<0.001; *** : p<0.0001 2-way ANOVA followed by Sidak’s multiple comparison test.

## Additional supplementary file

**Supplementary Video 1 | Exemplary video showing mature swarming worms, as well as worms just before swarming**

## Notes

### Competing Interest Statement

The authors have declared no competing interest.

